# Acute IFN-γ elicits long-term proteomic IFN-γ signature in melanoma cells

**DOI:** 10.64898/2025.12.23.696219

**Authors:** Agnete W. P. Jensen, Anne-Christine Kiel Rasmussen, Ilaria Piga, Anne Weiss, Marta Sanchez Sanchez, Marco Donia, Giulia Franciosa, Jesper V. Olsen

**Affiliations:** Novo Nordisk Foundation Center for Protein Research, Department of Cellular and Molecular Medicine, Faculty of Health and Medical Sciences, Copenhagen University, Copenhagen, Denmark; National Center of Cancer Immune Therapy, Department of Oncology, Copenhagen University Hospital - Herlev and Gentofte, Herlev, Denmark; Samplix ApS, Birkerød, Denmark

**Keywords:** Melanoma cells, IFN-γ signaling, Mass spectrometry, Proteomics

## Abstract

Interferon-γ (IFN-γ) is a central mediator of antitumor immunity, and its downstream effects are widely assumed to require continuous cytokine exposure. Here, we applied state-of-the-art mass spectrometry (MS)-based proteomics to patient-derived melanoma cells exposed either continuously to IFN-γ or to a 3-hour pulse followed by cytokine removal. A transient IFN-γ exposure was sufficient to establish a stable proteomic state indistinguishable from continuous stimulation, demonstrating that IFN-γ-induced remodeling is rapidly triggered and durably maintained independently of ligand persistence. Proximal JAK-STAT signaling was rapid, saturable at low cytokine concentrations, and fully reversible upon washout, yet downstream proteomic programs remained durably imprinted. In contrast, functional consequences such as cytostasis remained dose-dependent, revealing a disconnect between signaling activation and phenotypic sensitivity. Together, these findings challenge long-standing assumptions about IFN-γ signaling, redefine the temporal requirements for cytokine-driven tumor cell programming, and provide a mechanistic framework for improving immunotherapies primarily working via IFN-γ-signaling and physiologically relevant *in vitro* models of tumor-immune interactions.

## Introduction

Interferon-gamma (IFN-γ) is a central immunoregulatory cytokine produced predominantly by activated T and natural killer (NK) cells, playing a pivotal role in antitumor immunity^1,2^. Engagement of IFN-γ with its receptor (IFNGR1/IFNGR2) on tumor cell surface triggers the canonical Janus kinase (JAK) and signal transducer and activator of transcription (STAT) pathway: The cytoplasmic tyrosine-protein kinases JAK1 and JAK2 are recruited and activated, leading to STAT1 phosphorylation, dimerization, and nuclear translocation^3^. This initiates a primary and secondary transcriptional response, inducing hundreds of interferon-stimulated genes (ISGs) that regulate antigen processing and presentation through upregulation of human leukocyte antigen (HLA) molecules, chemokine secretion (CXCL9/10/11), apoptosis, chromatin and metabolic remodeling, and immune checkpoint expression such as programmed cell death 1 ligand 1 (PD-L1)^2^.

A T cell-inflamed IFN-γ response signature has been shown to positively correlate with immune checkpoint inhibitor (ICI) response in patients with melanoma^4^. However, tumor cells frequently evade IFN-γ-mediated immune pressure through genetic and non-genetic mechanisms. Loss-of-function mutations in JAK1, JAK2, or IFNGR1/2 are well-documented contributors to ICI resistance in melanoma^5–8^, and additional genetic alterations, such as amplification of the negative regulator SOCS1, can further dampen IFN-γ signaling^5^. However, the majority of ICI-resistant tumors harbor no such mutations^9^. Beyond JAK-STAT signaling, IFN-γ can engage auxiliary pathways, including PI3K-AKT (shown in lung adenocarcinoma), where it can promote proliferation and survival^10,11^, thereby contributing to context-dependent modulation of tumor cell fate. In addition to well-known inhibitors like SOCS proteins, more recently described negative regulators, including LNK and STUB1, have also been shown to dampen IFN-γ responses in melanoma cells^12,13^. While these adaptations underscore the importance of IFN-γ as a selective pressure in the tumor microenvironment (TME), they also emphasize the need to better understand how IFN-γ signaling is initiated, sustained, and maintained at the cellular level.

*In vitro* studies routinely expose tumor cells to continuous IFN-γ for 48-72 hours, implicitly assuming that sustained stimulation is necessary to achieve maximal signaling and cellular responses^14–16^. Despite decades of IFN-γ research, a fundamental question remains largely unexplored: how long must a tumor cell “see” IFN-γ to commit to a stable IFN-γ-induced response program? Is a brief, physiologically relevant pulse sufficient to trigger a durable transcriptional and proteomic program, or is continuous exposure required? Addressing this question is essential for establishing physiologically meaningful *in vitro* models and understanding how T cells impose selective pressure on tumors during natural immune surveillance.

To date, proteomic studies of IFN-γ signaling have relied on immortalized cell lines (A375 melanoma, A549 lung carcinoma, HEK293T, or endothelial cells) using data-dependent acquisition (DDA) and continuous stimulation paradigms^17–19^. Comprehensive proteomic maps of IFN-γ-treated primary cancer cells remain absent. In this study, we address this gap by performing state-of-the-art, data-independent acquisition (DIA) mass spectrometry on primary melanoma cell lines treated with either continuous IFN-γ or a transient 3-hour pulse followed by cytokine washout. This approach enables us to define the temporal requirements for IFN-γ-induced cellular reprogramming and uncover the durability of downstream proteomic responses.

## Methods

### Sample origin, establishment of primary melanoma cell lines and autologous REP TILs

All procedures were approved by the Scientific Ethics Committee for the Capital Region of Denmark. Tumor biopsies were obtained from three patients diagnosed with cutaneous metastatic melanoma who were enrolled in the clinical trials at the National Center for Cancer Immune Therapy (CCIT-DK), Department of Oncology, Copenhagen University Hospital, Herlev, Denmark (H-20070020). Written informed consent was obtained from all patients prior to sample collection, in accordance with the principles of the Declaration of Helsinki.

Fresh tumor specimens were transported from the clinic to the laboratory in RPMI 1640 medium (Gibco, Thermo Fisher Scientific). Upon arrival, samples were dissected to remove surrounding non-tumor tissue and mechanically dissociated into 1-3 mm³ fragments. Primary melanoma cell lines were established by short-term *in vitro* culture and serial passaging of adherent tumor cells in complete RPMI medium under standard culture conditions (37 °C, 5% CO₂). Established cell lines were authenticated based on morphological features, growth characteristics, and expression of melanoma-associated markers, as assessed by flow cytometry (described below).

For the tumor cell line (TCL)24, autologous tumor-infiltrating lymphocytes (TILs) were isolated from the same tumor biopsy and expanded *ex vivo* using a two-step protocol as previously described^20^. Briefly, minimally cultured “young” TILs were generated, followed by large-scale expansion using a rapid expansion protocol (REP).

### Cell culture and IFN-γ stimulation

Primary melanoma cell lines were maintained at 37 °C in a humidified incubator with 5% CO₂. Cells were cultured in RPMI 1640 Medium with GlutaMAX (Gibco, Thermo Fisher Scientific), supplemented with 25 mM HEPES (Gibco, Thermo Fisher Scientific), 10% fetal bovine serum (FBS, Gibco, Thermo Fisher Scientific), and 100 U/mL penicillin and 100 µg/mL streptomycin (Pen-Strep, Gibco, Thermo Fisher Scientific). Cells were routinely tested for mycoplasma contamination by PCR (Sartorius) following the manufacturer’s instructions. No Mycoplasma infection was detected in the past four years.

For stimulation experiments, cells were seeded and, 24 hours later, “starved” overnight in a culture medium containing 1% FBS to reduce basal signaling. Subsequently, cells were treated with recombinant human interferon-γ (IFN-γ, PeproTech) at the indicated concentrations and durations depending on the downstream application. Short-term stimulations (30 minutes to 6 hours) were used for phosphorylation analyses, including Western blotting and phosphoproteomics, whereas long-term stimulations (24 hours to 72 hours) were performed for proteomic profiling and cell viability assays. For all proteomic experiments, a time-matched untreated control was included. To assess the dose- and time-dependence of IFN-γ signaling, melanoma cells were exposed to varying concentrations of IFN-γ (0-800 IU/mL) for the indicated durations. To evaluate signaling reversibility, washout experiments were conducted in which cells were stimulated with 50 IU/mL IFN-γ for 3 hours, 3+3 hours (IFN-γ renewed after 3 hours), or 6 hours, followed by replacement of the medium with fresh IFN-γ-free medium until harvest at 48 hours. In selected experiments, cells were stimulated with IFN-γ in the presence or absence of protease and phosphatase inhibitors (Protease Inhibitor Cocktail, Sigma-Aldrich). At harvest time points, cells were washed twice with Dulbecco’s phosphate-buffered saline (PBS, Sigma-Aldrich/Merck KGaA) and processed for downstream analyses.

### Real-time viability analysis using xCELLigence

Real-time monitoring of tumor cell viability following IFN-γ stimulation was performed using the xCELLigence RTCA eSight™ system (Agilent). Primary melanoma cells were seeded into 96-well RTCA E-plates (Agilent) in complete RPMI 1640 medium at densities optimized for each cell line. To assess long term effect on growth and viability, cells were seeded at low density to allow growth for up to 5 days; TCL05 and TCL15 at 10.000 cells/well and TCL24 at 5.000 cells/well in 200 ul media. After 24 hours, 100 ul media was removed at replaced with 100 µL fresh medium containing IFN-γ (range 0-200 IU/mL) and the measurements were resumed. Cell Index (CI) values were recorded every 15 minutes until 87 hour post IFN-y treatment CI values were normalized to the last measurement immediately prior to IFN-γ addition (normalized CI, NCI), and changes in NCI were tracked over time. To evaluate the impact of IFN-γ pretreatment on tumor cell killing by autologous TILs, TCL24 cells were pretreated with IFN-γ for 72 h (range 0-200 IU/mL), harvested, and re-seeded in the xCELLigence system at 10,000 cells/well in 200 µL medium to achieve a CI of approximately 1 after 24 h. After 24 h, 100 µL medium was removed and replaced with 100 µL fresh medium containing TILs, or no effector cells (tumor-only control), at an effector-to-target ratio of 1:1 (TIL:TCL). CI values were then recorded continuously for up to 120 h following effector cell addition. CI was normalized to 1 at the time point immediately prior to effector cell addition, and relative killing (%) was calculated as: 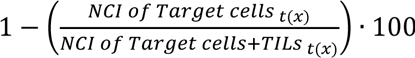. Each experimental condition was performed in two technical replicates.

### Flow cytometry

Flow cytometry was used for both phenotypic characterization of primary melanoma cell lines and for assessing IFN-γ-induced changes in surface marker expression over time.

For characterization, 5×10^5^ cells were harvested and washed twice with PBS. Cells were stained with Live/Dead™ Fixable Near-IR viability dye (APC-Cy7, Thermo Fisher Scientific) and the following antibodies: anti-MCSP (PE, clone EP-1, Miltenyi Biotec), anti-CD146 (BV421, clone P1H12, BioLegend), and anti-CD90 (FITC, clone 5R10, BioLegend). Staining was performed in PBS containing 0.1% FBS at 4 °C for 30 minutes in the dark. After staining, cells were washed twice with PBS and analyzed on a NovoCyte Quanteon™ Flow Cytometer (Agilent).

For IFN-γ experiments, melanoma cells were treated as described in the “Cell culture and IFN-γ stimulation” section. At each time point, cells were harvested, washed, and stained with Live/Dead™ viability dye and antibodies against MHC class I (HLA-ABC, APC, clone G46-2.6, BD Biosciences), MHC class II (HLA-DR/DP/DQ, FITC, clone TÜ39, BD Biosciences), and IFN-γ receptor 1 (IFN-γR1/CD119, PE, clone GIR-208, Biolegend). Isotype controls were included for MHC staining. Staining and acquisition were performed as described above.

Flow cytometry data were analyzed using NovoExpress software version 1.5.6 (Agilent). Gates were defined based on unstained and isotype control samples.

### Enzyme linked immunosorbent assays (ELISA)

The concentration of IFN-γ in cryopreserved culture supernatants was quantified using ELISA according to the manufacturer’s instructions. Absorbance was measured using a FLUOstar Omega (BMG Labtech).

### Western Blotting

Total protein content was analyzed using standard western blot protocols. Cells were seeded in 6-well plates at 2×10⁴ cells/well and cultured under the conditions described in the “Cell culture and IFN-γ stimulation” section. After 30 minutes of stimulation, cells were lysed in RIPA buffer (50 mM Tris-HCl, pH 7.5; 150 mM NaCl; 1% NP-40; 0.1% sodium deoxycholate; 1 mM EDTA; 5 mM β-glycerophosphate; 5 mM sodium fluoride; 1 mM sodium orthovanadate) supplemented with a protease inhibitor cocktail (cOmplete mini EASY pack, Merck). Lysates were incubated on ice for 30 minutes and centrifuged at 13,000 rpm for 15 minutes at 4°C. Supernatants were collected, and protein concentrations were determined using the Pierce BCA Protein Assay Kit (Thermo Fisher Scientific).

Equal amounts of protein were mixed with 4×NuPAGE LDS sample buffer (Thermo Fisher Scientific) supplemented with 100 mM dithiothreitol (DTT) and boiled at 70°C for 15 minutes. Proteins were separated by SDS-PAGE using Bis-Tris gels in MOPS buffer (Invitrogen) and transferred onto nitrocellulose membranes (Invitrogen) using the iBlot system (Thermo Fisher Scientific). Membranes were blocked for 1 hour at room temperature in 5% BSA in PBST (0.2% Tween 20 in PBS) and incubated overnight at 4°C with primary antibodies: rabbit anti-STAT1, rabbit anti-phospho-STAT1 (pTyr701, Cell Signaling Technology), and mouse anti-GAPDH, diluted 1:1000 in blocking buffer. Membranes were then washed with PBST and incubated for 1 hour at room temperature with HRP-conjugated secondary antibodies (1:10,000 in 5% milk in PBST), followed by additional washes. Protein detection was performed using the Novex™ ECL Chemiluminescent Substrate Reagent Kit (Thermo Fisher Scientific).

### Sample preparation for proteomic and phosphoproteomic analysis

#### Cell lysis and protein extraction

Cells were lysed directly in boiling SDS lysis buffer (5% sodium dodecyl sulfate (SDS), 100 mM Tris-HCl pH 8.5, 5 mM tris(2-carboxyethyl)phosphine (TCEP), 10 mM chloroacetamide (CAA)) and incubated for 10 minutes at 99°C while shaking. Lysates were subsequently sonicated, and protein concentrations were determined using the Pierce BCA Protein Assay Kit (Thermo Fisher Scientific).

#### Protein digestion

For each experiment, equal amounts of protein (88-350µg) were digested using the PAC (protein aggregation capture) protocol^21^. Briefly, proteins were aggregated on magnetic MagReSyn® Hydroxyl-microparticles (Resyn Biosciences) by addition of acetonitrile to 80% and proteins were digested on-bead overnight with Lys-C (FUJIFILM Wako Pure Chemical Corporation) and Trypsin (Sigma-Aldrich) at enzyme-to-protein ratios of 1:500 and 1:250, respectively, on the KingFisher Flex magnetic particle processing robot (Thermo Fisher Scientific). The following day, samples were acidified with trifluoroacetic acid (TFA) to a final concentration of 1%. For proteome analysis, either 0.075µg of peptides were loaded onto Evotips (Evosep Biosystems) for 40 SPD acquisition, or 0.5µg of peptides were injected using the Vanquish Neo system (Thermo Fisher Scientific) for 180 SPD acquisition. The remaining peptides (80µg) were desalted using C18 Sep-Pak cartridges (Waters Corporation) and eluted directly into plates for Zr-IMAC phosphopeptide enrichment.

#### Enrichment of phosphorylated peptides

The phosphopeptides were enriched by zirconium ion (Zr4+) chelated immobilized metal affinity chromatography (IMAC) performed on a KingFisher Flex robot (Thermo Fisher Scientific) in a 96-well format, as described previously^22,23^. Briefly, peptides were incubated with 8µL of magnetic Zr-IMAC HP beads (ReSyn Biosciences). Eluted phosphopeptides were acidified with 40 µL of 10% TFA and loaded onto Evotips (Evosep Biosystems) for LC-MS/MS analysis.

### LC-MS/MS analysis

Peptide samples were analyzed on an Orbitrap Astral mass spectrometer (Thermo Fisher Scientific)^24^ coupled to either an Evosep One LC system (Evosep Biosystems)^25^ or a Vanquish Neo UHPLC (Thermo Fisher Scientific), and interfaced online using an EASY-Spray ion source. Analytical column configurations were selected according to the experimental setup (**Supplementary Table 1)**. The instrument was operated in positive ion mode using narrow-window DIA^24^. The spray voltage was set to 1.8-2.0 kV, the capillary temperature to 275-280 °C, and the funnel RF level to 40. Full MS scans were acquired at a resolution of 180K or 240K with a scan range of *m/z* 380-980 for proteome analyses and *m/z* 480-1080 for phosphoproteome analyses. The automatic gain control (AGC) target was set to 500% with a maximum injection time of 3 or 30 ms. Precursor ions were fragmented by higher-energy collisional dissociation (HCD) with a normalized collision energy (NCE) of 25%. The fragment scan range was set at 150-2000 m/z. DIA-MS/MS fragment ion scans were recorded with a 2 or 4 Th quadrupole isolation window and a maximum injection time of 2.5, 5, or 6 ms. Detailed MS acquisition parameters for each experiment are provided in (**Supplementary Table 1)**.

### Raw mass spectrometry data processing

MS raw files were searched in Spectronaut version 20 (Biognosys) with a library-free approach (directDIA+) against the Human Uniprot fasta file (downloaded in October 2024 and contained 20,428 entries) and the cell culture contaminant fasta file (downloaded from https://github.com/HaoGroup-ProtContLib, with 370 entries)^26^.

All proteome data were searched together. Cysteine carbamylation was set as a fixed modification, and N-terminal acetylation and methionine oxidation were set as variable modifications and cross-run normalization was unchecked.

For phosphoproteomics, serine, threonine, and tyrosine phosphorylation were added as variable modifications. For phosphoproteomics analysis, the PTM localisation filter was checked, with a threshold of 0.75.

### Bioinformatic analysis

All bioinformatic analysis was performed using R version 4.5.2 with R studio version 2025.9.2.

#### Proteomics data filtering and processing

For all datasets, contaminants and entries without a gene name were removed, and MS intensities were log2 transformed. No imputation step was carried out.

The proteome dataset was split into experiment-specific tables for further analysis. For each proteomics dataset, intensities were normalized by median realignment. Principal component analysis (PCA) was conducted on experiment-specific datasets following exclusion of entries with missing values. For melanoma differentiation markers, normalized data were filtered to retain control samples without washout. Selected markers (MLANA, MITF, and PMEL) were mean-centered per protein and visualized as a heatmap.

For HLA analysis, normalized data were filtered for HLA-encoded proteins. Samples were annotated by TCL, treatment (control or IFN-γ), and washout status. Protein intensities were z-score normalized per tumor cell line and averaged across replicates for each condition prior to heatmap visualization.

The phosphosite table was exported from Spectronaut, using Spectronaut’s default filtering parameters and filtered for localisation probability > 0.75. MS intensities were log2 transformed and values < 1 were replaced with NA. Data was filtered for valid value, allowing to conserve phosphosites quantified in 50% of the runs, and normalized through median centering. No imputation step was carried out.

#### Differential expression analysis (DEA)

DEA was performed using the “limma” R package^27,28^.To account for repeated measurements from the same TCL, inter-sample correlation was estimated using the “duplicateCorrelation” function and incorporated into the linear model via the “block” argument in “lmFit”^29^. For analyses performed with limma alone, variance–intensity dependence was modeled using the trend = TRUE option. For DEqMS-based analysis, data were filtered by at least four of six replicates in at least one experimental group. Differential protein abundance was assessed using the DEqMS package^30^, which extends limma by modeling protein-level variance as a function of precursor counts. P-values were adjusted for multiple testing using the Benjamini-Hochberg procedure. Proteins were considered significantly regulated if they met both a statistical significance threshold (adjusted p-value ≤ 0.01) and a data-driven effect size threshold defined as an absolute log2 fold change exceeding two standard deviations of the corresponding logFC distribution (calculated separately for up- and down-regulated proteins). For analyses performed with limma alone, data were filtered by at least three of four replicates in at least one group. Volcano plots were generated using DEqMS- or limma-derived statistics, displaying log2 fold change versus −log10 adjusted p-value, with the top 20 most significantly up- and down-regulated proteins annotated

#### Single-sample gene set enrichment analysis (ssGSEA)

ssGSEA^31^ was performed on GenePattern using the standard settings.

#### STRING network analysis

Pathway enrichments were analyzed using STRING^32^.

#### Data visualization

Plots were performed using the R package “ggplot2” version 4.0.1 or the GraphPad Prism software version 10.

## Results

### Transient IFN-γ exposure induces a sustained proteomic response in melanoma cells

We first confirmed the absence of detectable fibroblast contamination and characterized differentiation heterogeneity among the melanoma cell lines (TCL05, TCL15, and TCL24) using three reliably detected markers (out of five commonly used^33^; **Figure S1A-C)**. We then investigated whether continuous IFN-γ stimulation is required to sustain the proteomic response by treating patient-derived melanoma cell lines either continuously for 48 hours or transiently for 3 hours followed by 45 hours in IFN-γ-free medium (washout). Whole-proteome profiling was performed at 48 hours across six biological replicates comparing the global protein expression changes induced by the two stimulation regimens **(Figure 1A)**, resulting in the identification of more than 5,600 protein groups on average across samples **(Figure S2A)**. Principal component analysis (PCA) revealed a near-complete overlap between the continuous and washout conditions across all three melanoma cell lines **(Figure 1B and S2B)**. Consistently, comparison of the significantly differentially expressed proteins revealed a strong correlation between the two conditions **(Figure 1C)**, indicating that the continuous and washout stimulation regimens elicit highly similar proteomic responses. To assess the concordance of IFN-γ-induced proteins between stimulation regimens, we compared volcano plots for IFN-γ versus control under continuous and washout conditions. Both analyses yielded an almost identical set of significantly regulated proteins, with the top 20 most significant hits, based on adjusted p-value and log₂ fold change, consisting of canonical ISGs, including STATs, TAP transporters, and HLA class I and II molecules **(Figure 1D and S2C)**. To further quantify pathway-level responses, we applied ssGSEA using IFN-γ-responsive gene sets from the Reactome and Hallmark collections within the Molecular Signatures Database (MSigDB)^34^. Across all three cell lines, ssGSEA scores for the IFN-γ signaling pathway increased markedly upon IFN-γ stimulation and were highly similar between the continuous and washout regimens **(Figure 1E and S2D)**, confirming equivalent activation of downstream IFN-γ signaling. Extraction of HLA protein abundances within the proteomics dataset revealed robust and comparable upregulation of multiple MHC class I and II components under both stimulation regimens **(Figure S2E)**. To independently validate these proteomic findings, flow cytometry confirmed sustained surface expression of classical IFN-γ-induced markers, including MHC class I (HLA-A, -B, -C), MHC class II (HLA-DR, -DP, -DQ), and PD-L1, across both continuous and washout treatments **(Figure 1F).** Collectively, these results demonstrate that a brief IFN-γ pulse is sufficient to initiate and sustain the full downstream proteomic program in melanoma cells, rendering prolonged cytokine exposure unnecessary for the establishment of a stable IFN-γ-induced state.

**Figure 1:**
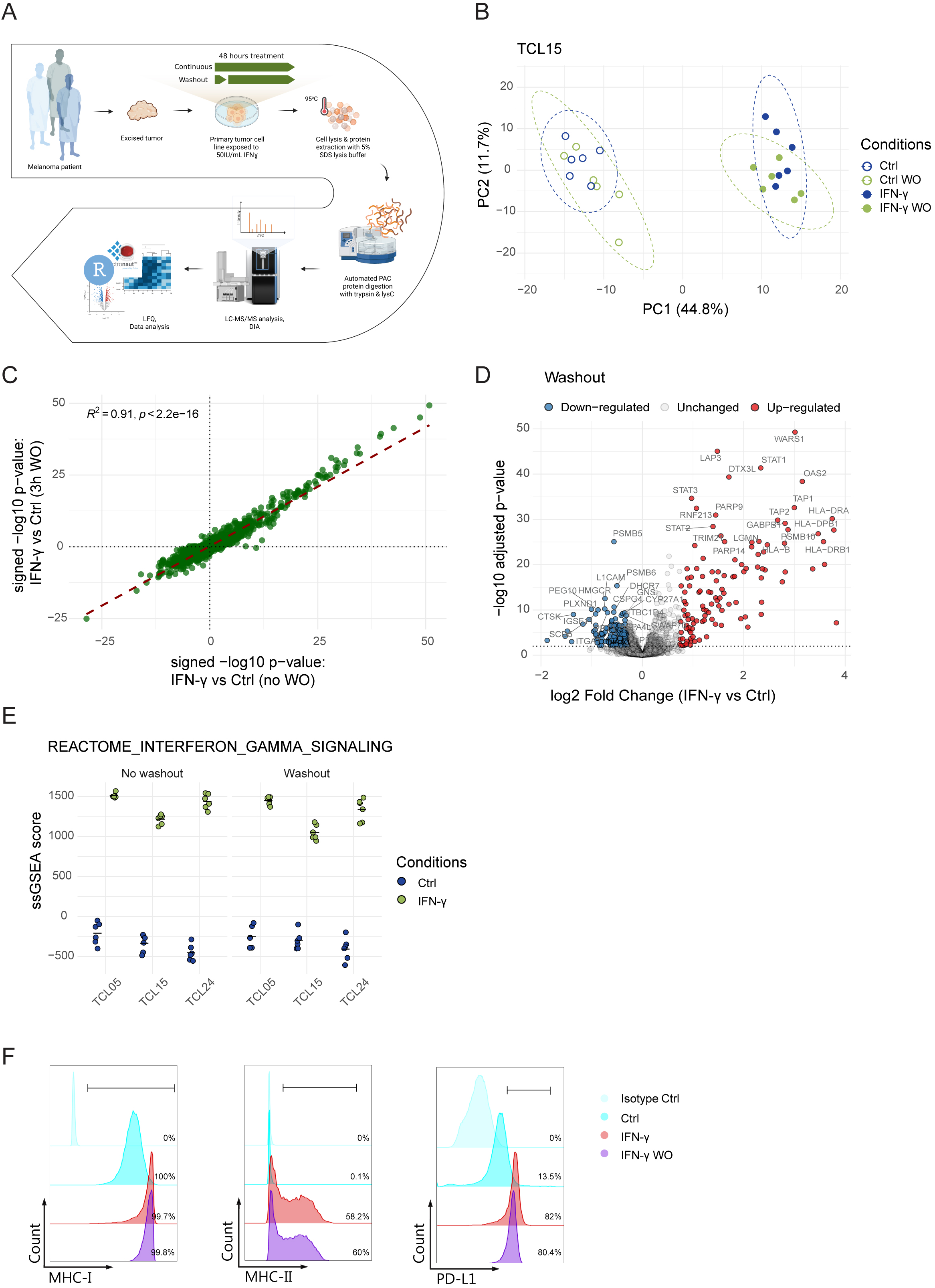
Continuous and transient IFN-γ exposure elicits equivalent proteomic changes in patient-derived melanoma cell lines. **(A)** Schematic drawing of the proteomics experiment design. Created with Biorender. **(B)** PCA of whole-proteome measurements from TCL15 cells subjected to continuous IFN-γ [50 IU/mL] stimulation for 48 hours (IFN-γ), transient stimulation for 3 hours followed by 45 hours in cytokine-free medium (IFN-γ WO), or matched control conditions (Ctrl and Ctrl WO). Ellipses show 95% confidence intervals for each condition. **(C)** Scatter plot showing the signed -log10 adjusted p-values for differential protein expression (IFN-γ vs control) under continuous stimulation (x-axis) and washout (WO) condition (y-axis). The dashed line represents the linear regression fit. **(D)** Volcano plot of differentially expressed proteins between the IFN-γ and control under washout conditions. DEqMS-derived, Benjamini-Hochberg FDR-adjusted p-values are shown. The top 20 up- and down-regulated proteins are annotated. **(E)** Single-sample gene set enrichment analysis (ssGSEA) using the REACTOME IFN-γ signaling pathway. **(F)** Histograms showing surface expression of MHC class I, MHC class II, and PD-L1 on TCL24 following the two stimulation regimes, including untreated controls and isotype controls, measured by flow cytometry (n=1).

### IFN-γ stability and receptor availability do not account for the sustained proteomic response

To determine whether ligand degradation or receptor internalization could account for the similar proteomic responses observed after transient and continuous IFN-γ exposure, we first assessed cytokine stability in the culture medium over time. TCL24 cells were cultured with or without IFN-γ, and supernatants were collected from 30 minutes to 48 hours for quantification by ELISA. Compared with the positive control, IFN-γ concentrations decreased sharply within the first 30 minutes after addition to the cells. This could be due to rapid ligand uptake and/or partial degradation by secreted proteases. Nonetheless, IFN-γ remained readily detectable with minimal further decay across all measured time points **(Figure 2A)**. As expected, IFN-γ was nearly undetectable in negative control samples. These measurements indicate that IFN-γ persists at stable, lower concentrations in the culture environment throughout the duration of the proteomics experiment. We next examined IFN-γ receptor availability at the cell surface by flow cytometry. IFNGR1 expression decreased within one hour of IFN-γ stimulation, however, this reduction was transient, as surface IFNGR1 levels returned to baseline by two hours, indicating rapid recycling to the plasma membrane **(Figure 2B)**. Given the reduced IFN-γ concentration after the initial exposure, we next evaluated whether providing additional ligand during the window in which IFNGR1 returns to the cell surface would alter the downstream proteomic response. TCL24 cells were stimulated with IFN-γ for three hours, replenished with fresh IFN-γ, and stimulated for an additional three hours, or stimulated continuously for six hours followed by washout. This design ensured that ligand was available during the period when IFNGR1 had recycled back to the plasma membrane, thereby supporting continued receptor engagement throughout the early phase of signaling. After the six-hour stimulation period, medium was replaced with cytokine-free medium, and cells were harvested at 48 hours for proteomic analysis. Consistent with our previous findings, no differences were observed between the pulse and continuous stimulation regimens, even when IFN-γ was reintroduced to provide an additional stimulus **(Figure 2C)**.

**Figure 2:**
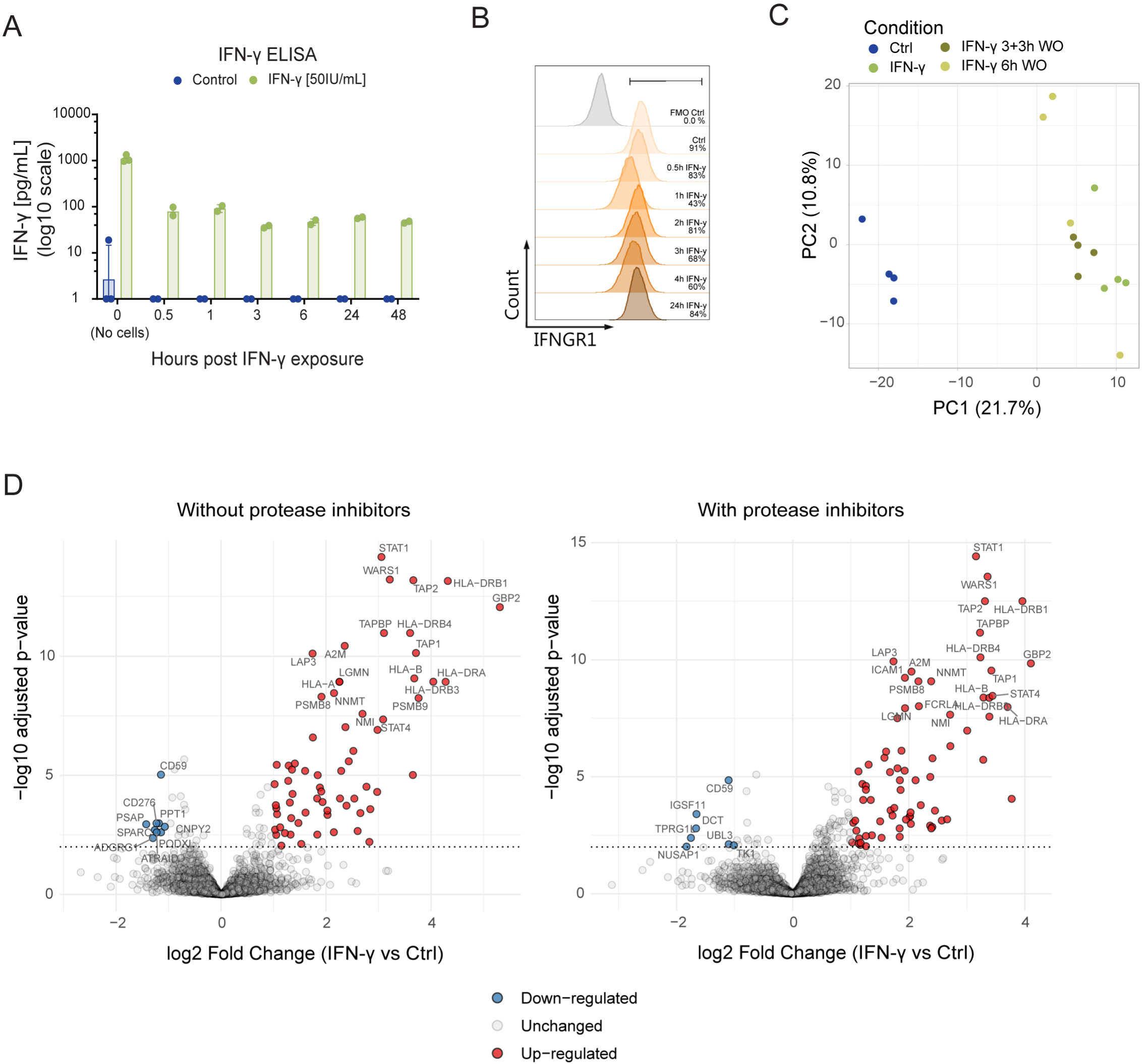
IFN-γ ligand stability, receptor dynamics, and protease activity do not account for the equivalence of continuous and transient stimulation. **(A)** ELISA-based quantification of IFN-γ in supernatants collected from TCL24 cells cultured with or without IFN-γ [50 IU/mL] from 30 minutes to 48 hours after cytokine addition (n=2). **(B)** Time-course flow cytometry analysis of IFNGR1 surface expression on TCL24 cells following IFN-γ stimulation [25 IU/mL] from 30 minutes to 24 hours (n=1). **(C)** PCA of TCL24. Cells were stimulated with IFN-γ [50 IU/mL] under modified dosing regimens. Cells were stimulated for 3 hours and either replenished with fresh IFN-γ for an additional 3 hours (3+3h WO) or stimulated continuously for 6 hours prior to washout (6h WO) (n=4 replicates per condition). **(D)** Volcano plot of differentially expressed proteins in TCL24 treated with IFN-γ compared to control at 48 hours in the absence (left) or presence (right) of protease inhibitors (PI). The top 20 up- and down-regulated proteins are annotated (n=4 replicates per condition).

To assess whether proteolytic activity during cell stimulation could cleave or inactivate IFN-γ, TCL24 cells were stimulated with IFN-γ in the presence or absence of broad-spectrum protease inhibitors (PI) and analyzed by MS after 48 hours. Across all conditions, more than 5,500 proteins were quantified **(Figure S3A)**. PCA revealed that IFN-γ stimulation was the primary driver of variance in the dataset, while PI treatment accounted for variation along the second principal component **(Figure S3B)**. Despite this separation, canonical IFN-γ-responsive proteins exhibited highly similar induction patterns with or without PI, indicating that extracellular proteolytic activity does not substantially attenuate IFN-γ signaling **(Figure 2D)**. Given the PCA-based separation between PI-treated and untreated samples, we next performed KEGG and subcellular localization enrichment analyses on proteins significantly altered by PI treatment. These analyses showed enrichment of pathways and compartments related to lysosomes, vesicles, and membrane-associated structures, including lysosome, lytic vacuole, vesicle, and bounding membrane of organelle terms **(Figure S3C-D)**. Accordingly, PI treatment led to a significant increase in IFNGR1 abundance **(Figure S3E-F)**, indicating receptor stabilization under conditions of reduced proteolysis. However, PI treatment did not cause a global increase in the abundance of membrane-associated proteins, indicating that PI does not broadly affect the stabilization of membrane proteins. Despite the increased availability of IFNGR1 during IFN-γ stimulation, downstream IFN-γ-responsive proteins remained unaffected **(Figure 2C)**, demonstrating that protease inhibition does not alter the canonical IFN-γ signaling output.

Collectively, these data demonstrate that the IFN-γ-dependent proteomic program is not limited by ligand degradation, extracellular proteolytic activity, or complete receptor downregulation. Instead, the results indicate that early receptor engagement triggers a self-sustaining, cell-intrinsic signaling cascade, establishing a lasting cellular state that persists at least 45 hours after transient IFN-γ exposure.

### Rapid saturation of proximal IFN-γ signaling contrasts with dose-dependent tumor-intrinsic and T cell-mediated killing

To determine how proximal IFN-γ signaling dynamics relate to the durability of the downstream proteomic program, we first performed temporal phosphoproteomic profiling of STAT1 activation across all three melanoma cell lines. IFN-γ stimulation induced a rapid and robust increase in STAT1 S727 phosphorylation within 30 minutes, a modification associated with maximal transcriptional activity^35^, which was sustained only under continuous ligand exposure. Upon ligand withdrawal (washout), pSTAT1 levels declined back toward baseline, confirming that proximal IFN-γ signaling is strictly ligand-dependent and intrinsically transient **(Figure 3A)**. To further assess if STAT1 phosphorylation was dose-dependent, TCL24 cells were stimulated with a serial dilution of IFN-γ for 30 minutes. STAT1 Y701 phosphorylation, the site required for STAT1 dimerization and nuclear translocation^35^, saturated at very low IFN-γ concentrations, with no further increase at higher doses **(Figure 3B)**. Despite this early saturation of proximal signaling, downstream functional responses exhibited marked dose dependence. Real-time impedance monitoring (xCELLigence) revealed a dose-dependent reduction in tumor cell growth across all three melanoma lines, although with substantial variation in sensitivity. TCL15 exhibited marked growth suppression at even moderate IFN-γ concentrations, whereas TCL05 and TCL24 showed only partial sensitivity and failed to reach complete growth arrest even at the highest doses tested **(Figure 3C)**. Notably, these functional differences emerged despite saturation of STAT1 phosphorylation at very low IFN-γ concentrations, indicating that ligand dose continues to modulate cellular responses independently of proximal STAT1 activation. To evaluate how IFN-γ dose shapes tumor cell susceptibility to T cell-mediated killing, TCL24 were pre-incubated with a serial dilution of IFN-γ prior to 2D co-culture with autologous TILs. Pre-exposure to IFN-γ markedly enhanced tumor cell killing in a dose-dependent manner, with higher IFN-γ concentrations resulting in substantially greater cytotoxicity **(Figure 3D)**. Together, these results demonstrate that, although proximal STAT1 signaling rapidly saturates, the downstream tumor-intrinsic and T cell-mediated consequences of IFN-γ exposure remain strongly dose dependent.

**Figure 3:**
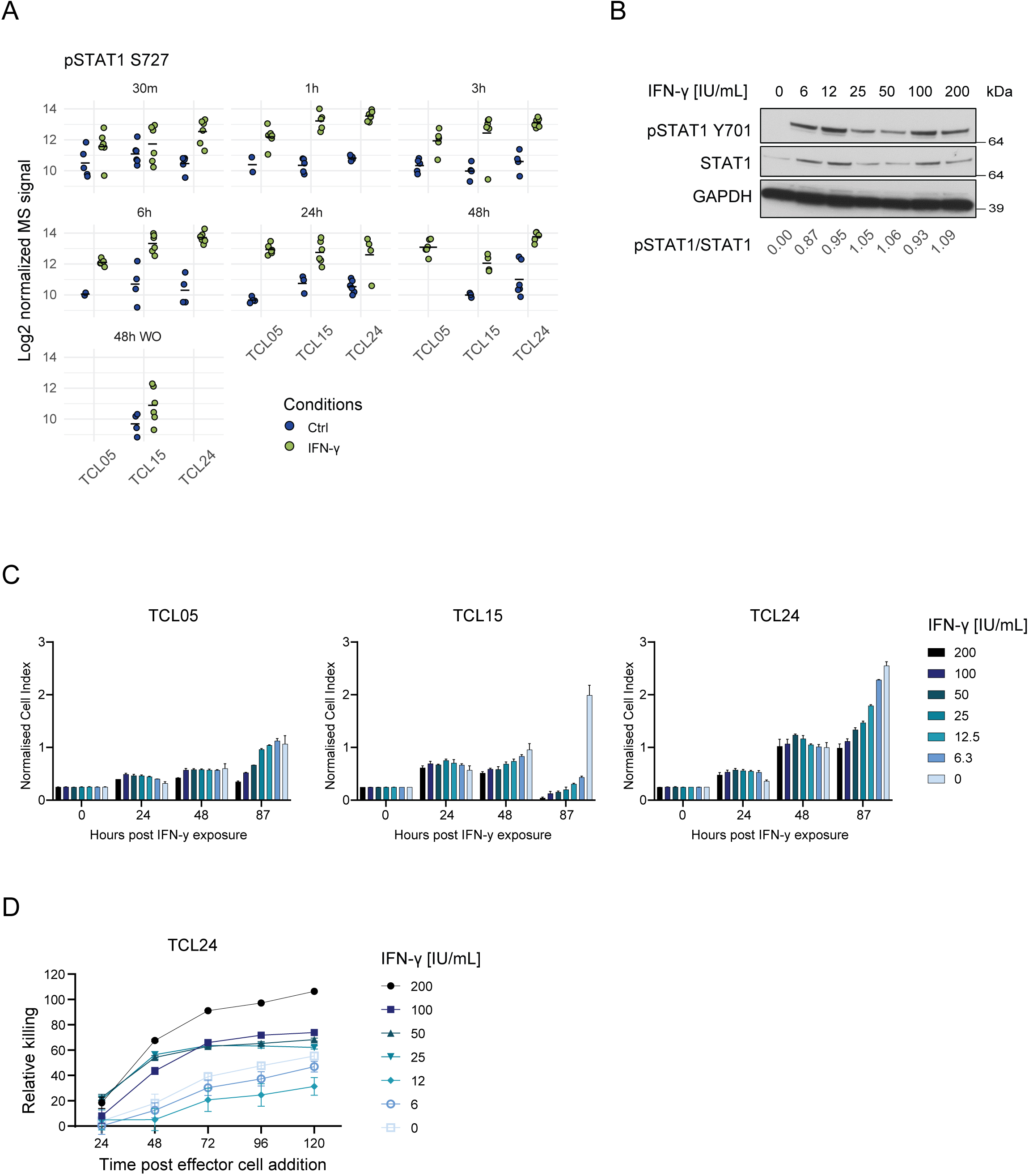
Functional responses to IFN-γ scale with dose despite saturating STAT1 phosphorylation. **(A)** Log2 MS intensities of STAT1 S727 phosphorylation from 30 minutes to 48 hours following IFN-γ [50 IU/mL] stimulation, including the washout condition (WO), across all three TCLs. The black dash represents the mean (n=6 replicates per condition per TCL). **B)** Western blot showing phosphorylation of STAT1 (pSTAT1 Y701), total STAT1, and GAPDH (loading control) in TCL24 stimulated for 30 minutes with increasing concentrations of IFN-γ [0-200 IU/mL] (n=1). **(C)** IFN-γ-mediated growth inhibition of TCL05, TCL15 and TCL24 measured by real-time impedance analysis (xCELLigence). Cells were stimulated with a serial dilution of IFN-γ [0-200 IU/mL] and growth assessed as the normalized Cell Index (NCI ± SD) (n = 2 technical replicates). **(D)** Autologous TIL-mediated killing of IFN-y pretreated TCL24 measured by xCELLigence at an effector-to-target (E:T) ratio of 1:1. Data are presented as mean ± SD (n = 2 technical replicates).

## Discussion

In this study, we demonstrate that a brief IFN-γ exposure in patient-derived melanoma cells is sufficient to induce a durable proteomic program that is indistinguishable from that elicited by continuous cytokine stimulation. While proximal IFN-γ signaling, assessed by STAT1 phosphorylation, was rapidly induced, saturated at low ligand concentrations, and fully reversible upon cytokine withdrawal, the downstream IFN-γ-responsive proteomic program persisted and remained strongly dose dependent. This uncoupling between transient upstream signaling and sustained downstream responses indicates that early IFN-γ pathway engagement is sufficient to establish a long-lasting cellular state and that IFN-γ dose sensing is not encoded solely at the level of STAT1 phosphorylation. One factor that may contribute to this behavior is IFN-γ-induced upregulation of STAT1. IFN-γ exposure markedly increased STAT1 abundance, thereby expanding the intracellular transcription factor pool. This increase could allow higher IFN-γ doses to drive greater cumulative transcriptional output over time. However, recent work demonstrates that STAT1 abundance is not rate limiting for the establishment or maintenance of IFN-γ-induced transcriptional memory. Instead, STAT1 activity is required during the initial priming phase to establish a durable cellular state, after which maintenance of this state becomes largely STAT1 independent and is not driven by continued target gene transcription^36^. These findings support a model in which early STAT1 engagement initiates IFN-γ-induced priming, whereas the persistence and dose dependency of downstream responses are encoded by regulatory mechanisms acting downstream of, or in parallel to, STAT1 activation. Such mechanisms are likely to include chromatin remodeling and epigenetic priming at ISG loci^33,37^, which may enable sustained responsiveness despite transient proximal signaling and early saturation of STAT1 phosphorylation. In addition, STAT1-independent IFN-γ-responsive pathways^38^ may further contribute to graded proteomic outputs as a function of cytokine dose and exposure duration.

We systematically excluded conventional explanations for the durability of downstream responses. Neither cytokine depletion, receptor downregulation, nor extracellular proteolysis accounted for the similarity between transient and continuous stimulation. IFNGR1 remained detectable at the cell surface throughout the time course, and broad-spectrum protease inhibition did not alter canonical IFN-γ-driven proteomic changes. Collectively, these findings support a model in which early receptor engagement is the dominant determinant of long-term cellular reprogramming. These findings are particularly relevant given how IFN-γ is delivered *in vivo*. Rather than persistent, uniform exposure, IFN-γ is released in transient, spatially restricted bursts at immune synapses, generating local cytokine gradients within the TME^39–41^. The near-identical proteomic programs induced by transient and continuous IFN-γ exposure in our system, especially across antigen-processing and presentation pathways, suggest that such brief cytokine pulses are sufficient to establish stable, immune-sensitized tumor cell states that preserve susceptibility to CD8⁺ T cell-mediated cytotoxicity^41^.

Our work also highlights key gaps for future investigation. Because proteomic measurements were only performed at the 48-hour time point, the kinetics of IFN-γ–induced proteome remodeling at earlier stages remain unresolved, leaving open questions about the temporal sequence of proteomic changes and intermediate signaling events. Similarly, we currently lack information on how the proteomic state evolves beyond 48 hours after cytokine removal, specifically if, when and how tumor cells begin to downregulate IFN-γ-induced proteins and return toward their basal state. While our data capture broad IFN-γ-induced proteomic remodeling, very low-abundance regulatory proteins such as SOCS1 and SOCS3 (key negative regulators of JAK-STAT signaling), remained below the detection threshold of our current dataset. These molecules may nonetheless play important roles in fine-tuning the magnitude and duration of IFN-γ responses, and their absence highlights the challenge of comprehensively quantifying low abounded proteins in global proteomic workflows. Addressing these gaps will be essential to fully understand the durability, reversibility, and regulatory constraints of IFN-γ-driven tumor cell reprogramming.

Collectively, our findings redefine the temporal requirements for IFN-γ-mediated tumor cell reprogramming. We show that a brief cytokine encounter is sufficient to lock melanoma cells into a stable, immune-altered state, challenging long-held assumptions about the duration of cytokine exposure required for tumor reprogramming. These insights have important implications for *in vitro* experimental design, and lay the groundwork for exploiting T cell-derived cytokine dynamics to improve anti-tumor immunity in solid tumors.

## Data Availability

The datasets generated and analyzed during the current study are available from the corresponding author upon request, subject to a collaboration agreement.

## Acknowledgments

We are grateful to all patients who donated their samples for this work.

## Authors’ contribution

GF and JVO designed the study. AWPJ wrote the original draft of the manuscript. AWPJ and ACKR generated the figures. AWPJ, ACKR, IP, AW, and MSS performed experiments. AWPJ, ACKR, and GF analyzed data. AWPJ and GF generated the R code used to analyze the data. JVO and MD provided resources and coordinated the project. GF, MD and JVO supervised the project and acquired funds. All co-authors read and edited the manuscript.

## Declaration of generative AI and AI-assisted technologies in the writing process

During the preparation of this work, the authors used ChatGPT version 5.2 in order to improve language and readability. After using this tool, the authors reviewed and edited the content as needed and take full responsibility for the content of the publication.

---

**Figure S1:**
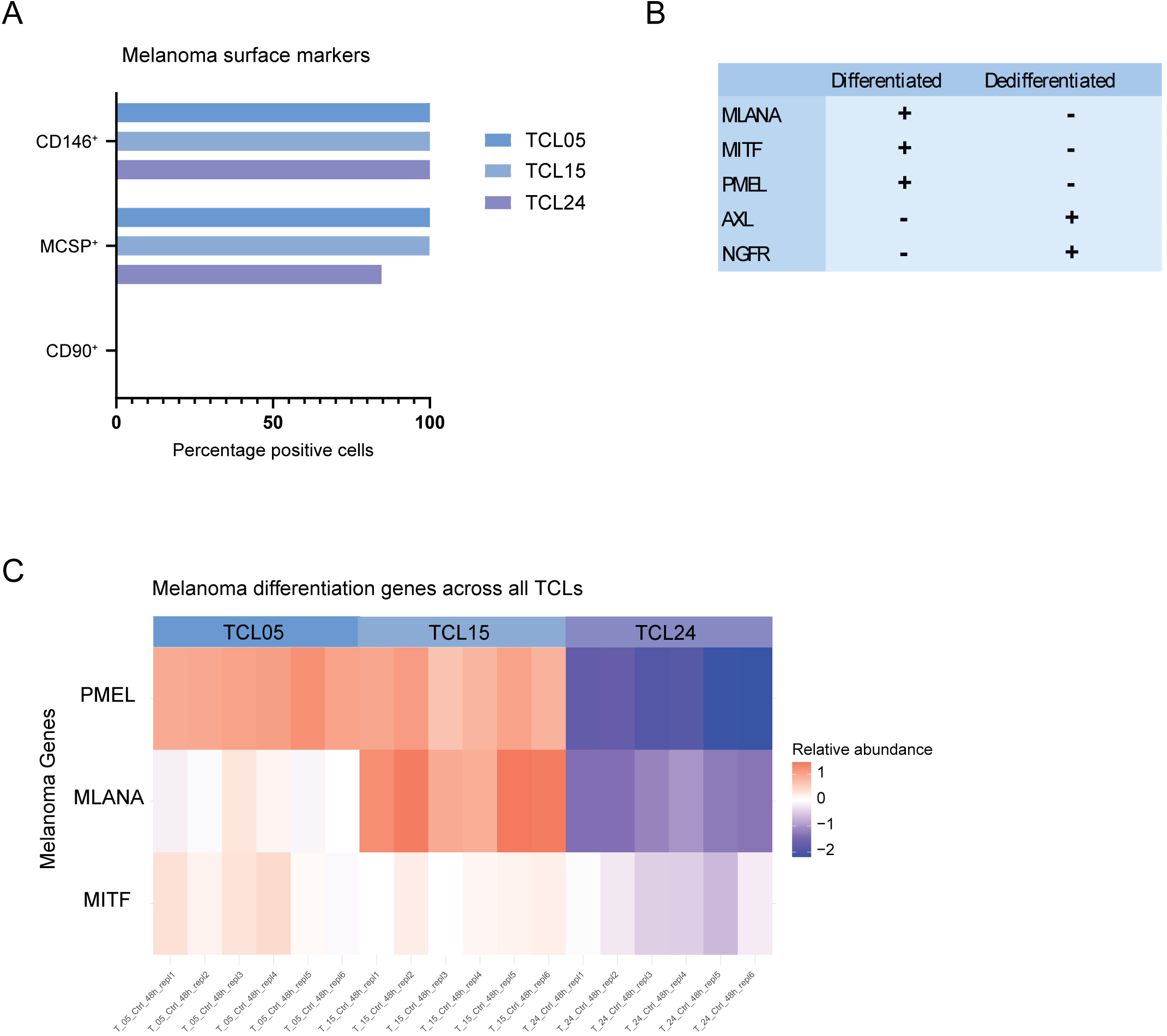
Characterization of primary melanoma tumor cell lines (TCLs). **(A)** Surface expression of two melanoma markers (CD146 and MCSP) and one fibroblast marker (CD90) in three different melanoma patient-derived TCLs. **(B)** Table summarizing melanoma lineage markers associated with differentiated (MLANA, MITF, PMEL) and dedifferentiated (AXL, NGFR) phenotypes^33^. **(C)** Heatmap of relative expression of melanoma differentiation markers (PMEL, MLANA, and MITF) under control conditions across three TCLs.

**Figure S2:**
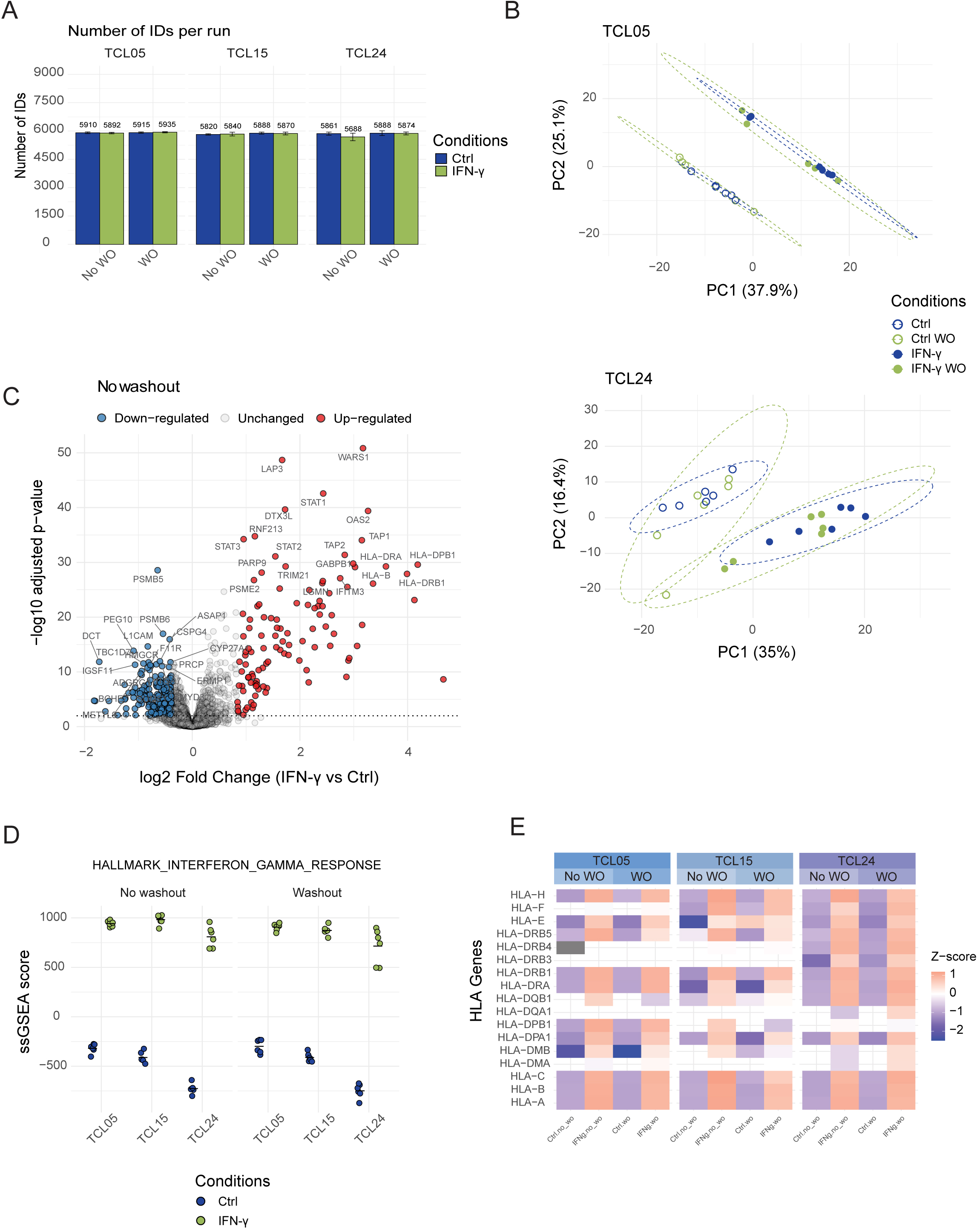
Supplementary analyses confirm that continuous and washout IFN-γ stimulation generate indistinguishable proteomic states. Related to Figure 1. **(A)** Protein group identifications across all three TCLs at 48 hours. Data is presented as mean ± SD (n=6 replicates per condition per TCL). Samples were analyzed on a 180 sample-per-day gradient. **(B)** PCA of whole-proteome measurements from TCL05 (top) and TCL24 (bottom). See additional description of experiment in Figure 1B. **(C)** Volcano plot of differentially expressed proteins between the IFN-γ [50 IU/mL] and control. DEqMS-derived, Benjamini-Hochberg FDR-adjusted p-values are shown. The top 20 up- and down-regulated proteins are annotated. See additional description of experiment in Figure 1D. **(D)** Single-sample gene set enrichment analysis (ssGSEA) using the Hallmark gene set for IFN-γ signaling pathway. **(E)** Heatmap of Z-scored log2 MS intensities of HLA class I and class II molecules in the three primary TCLs under control, continuous IFN-γ stimulation, or transient IFN-γ stimulation followed by washout.

**Figure S3:**
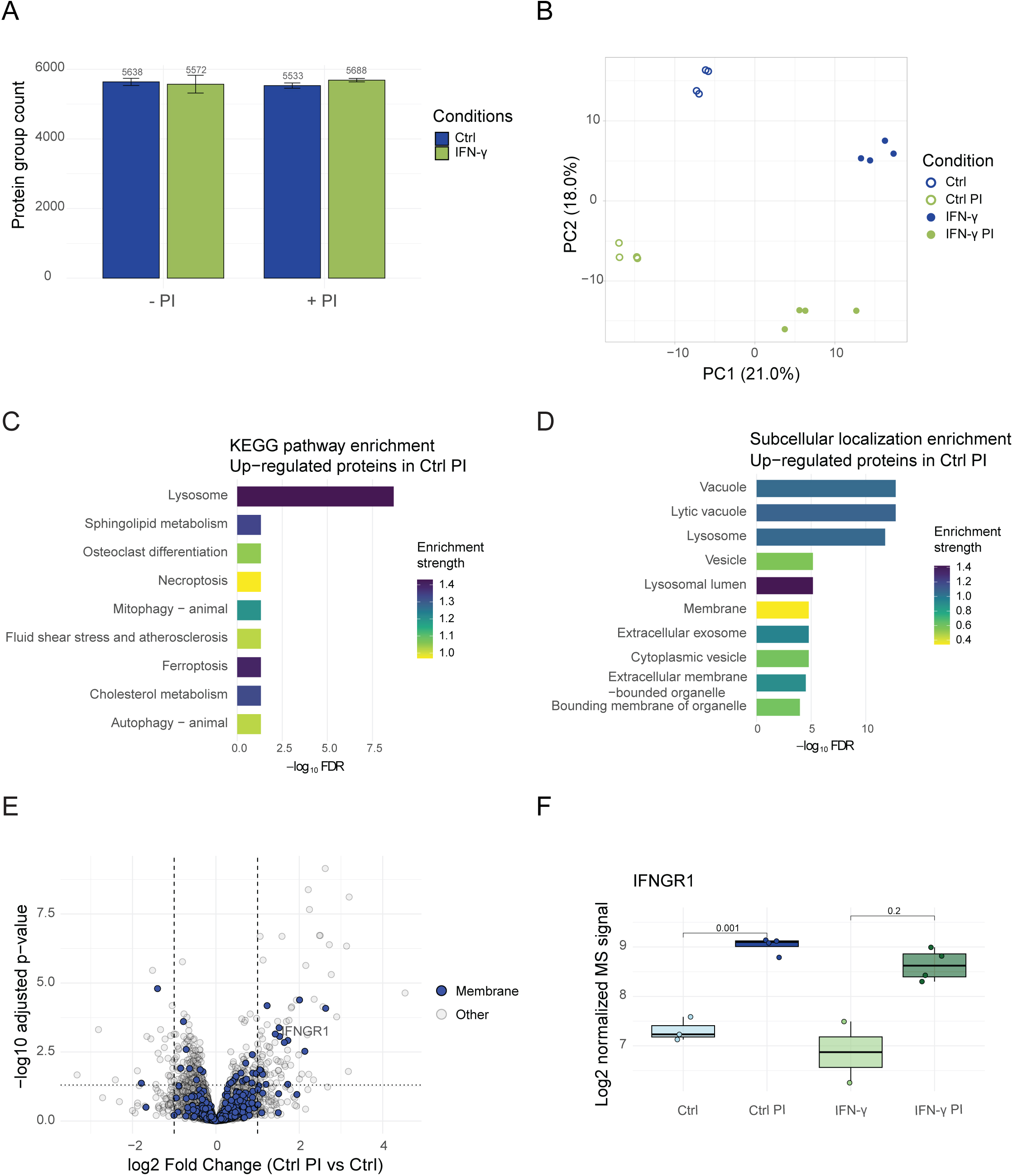
Protease inhibition does not alter IFN-γ-induced proteomic responses. Related to Figure 2. **(A)** Protein group identifications in TCL24 at 48 hours. Data are presented as mean ± SD (n=4 replicates per condition). Samples were analyzed on a 40 sample-per-day gradient. **(B)** PCA of TCL24. Cells were stimulated with IFN-γ [50 IU/mL] for 48 hours in the presence or absence of protease inhibitors (PI) (n=4 replicates per condition). **(C)** KEGG pathway and **(D)** subcellular localization (SCL) over-representation analyses performed using STRING for proteins significantly upregulated in TCL24 cultured in the presence of PI for 48 hours. A total of 59 entries were included in the analysis, and the most significantly enriched terms (lowest adjusted p-values) are shown (KEGG = all, SCL = Top10). **(D)** Volcano plot of differentially expressed proteins between the control and control with PI. Membrane proteins are marked in blue. **(E)** Log2 MS intensities of IFNGR1 at 48 hours. The black dash represents the mean (n=4 replicates per condition).

